# Noradrenergic tuning of arousal is coupled to coordinated movements

**DOI:** 10.1101/2024.06.18.599619

**Authors:** Li Li, Akshay N. Rana, Esther M. Li, Myesa O. Travis, Michael R. Bruchas

## Abstract

Matching arousal level to the motor activity of an animal is important for efficiently allocating cognitive resources and metabolic supply in response to behavioral demands, but how the brain coordinates changes in arousal and wakefulness in response to motor activity remains an unclear phenomenon. We hypothesized that the locus coeruleus (LC), as the primary source of cortical norepinephrine (NE) and promoter of cortical and sympathetic arousal, is well-positioned to mediate movement-arousal coupling. Here, using a combination of physiological recordings, fiber photometry, optogenetics, and behavioral tracking, we show that the LC^NE^ activation is tightly coupled to the return of organized movements during waking from an anesthetized state. Moreover, in an awake animal, movement initiations are coupled to LC^NE^ activation, while movement arrests, to LC^NE^ deactivation. We also report that LC^NE^ activity covaries with the depth of anesthesia and that LC^NE^ photoactivation leads to sympathetic activation, consistent with its role in mediating increased arousal. Together, these studies reveal a more nuanced, modulatory role that LC^NE^ plays in coordinating movement and arousal.

## 1 INTRODUCTION

As the integrated physiological responsiveness to sensory inputs, an animal’s arousal is closely matched to the needs of the animal’s behavior^1^. While arousal can be dissociated from movement such as in predatorprey contexts, initiating a movement is typically associated with needs for increased attention and metabolic consumption, whereas completing a movement is conversely associated with decreased needs^2–4^. How specific mechanisms in the brain coordinate movement and arousal remains unclear, and an active area of study. Understanding movement and arousal may provide new mechanistic insights into several neurodegenerative, psychiatric, and sleep pathologies. The locus coeruleus (LC) is a brainstem nucleus that sends noradrenergic projections across the cortex to regulate arousal states^5–7^, attention^8–10^, and sensory-motor functions^11–13^. Previous studies have shown LC activity to be absent during a cataplectic attack, but this loss of muscle tone can be partially restored by activating α_1_ adrenergic receptors (ARs)^14,15^. These findings suggest that the LC-norepinephrine (LC^NE^) system is important for movement-arousal coupling in healthy individuals.

Here, we report in a mouse model that during emergence from isoflurane (ISO) anesthesia, LC^NE^ activation is strongly coupled to the return of righting reflex (RORR), the moment when the animal regains organized movement. Similarly in awake mice, we observe characteristic patterns of LC^NE^ activation at movement initiations, and of LC^NE^ deactivation at movement arrests. Additionally, we find that LC^NE^ activity covaries with anesthetic depth, and that selective LC^NE^ activation increases sympathetic arousal, consistent with a role for LC^NE^ in arousal and wakefulness modulation. Combined, these results indicate that LC^NE^ neurons modulate their activity in response to movement to exquisitely tune arousal.

## 2 RESULTS

### Sequential neuronal activation during arousal transition

Arousal levels can range from absent in coma to high levels during active behavior^16,17^. While the LC^NE^ system is known for regulating sleepwake^18^ and stress^19^ which both fall within the middle of the arousal spectrum, its contribution at the two ends of the spectrum remains unclear. We first sought to address this question by measuring LC^NE^ activity in a controlled setting using emergence from anesthesia in a mouse model. This process nicely captures the transition from a pharmacologically induced coma to an awake behaving animal in a defined time window to measure physiological states using multiple measures. To this end, we expressed a cre-inducible fluorescent calcium sensor (AAV-DJEF1α-DIO-GCaMP6s) in the LC^NE^ neurons of DBH-cre mice (**Figure 1A**). We measured calcium dynamics of these neurons via fiber photometry while simultaneously recorded frontal electroencephalogram (EEG), heart rate, and behavioral movement (**Figure 1B**).

**FIGURE 1.**
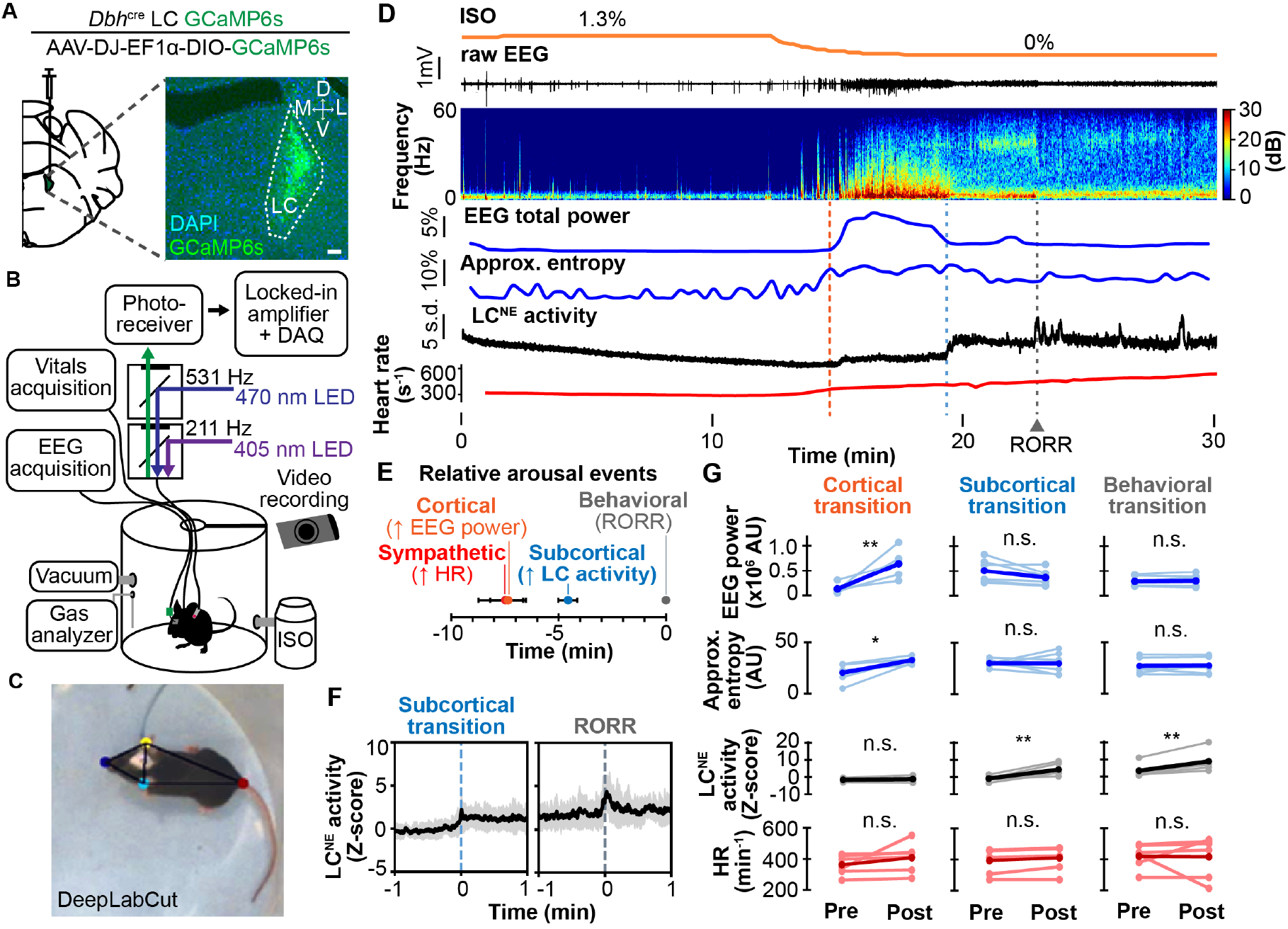
Locus coeruleus (LC) activity is robustly increased with movement at return of righting (RORR). **(A)** Left, schematic shows injection of GCaMP6s-expressing virus into the LC of Dbh-cre mice. Right, image shows GCaMP6s expression in the LC. Scale bar is 100 µm. **(B)** Diagram shows the anesthetic and recording setup for simultaneous acquisition of vitals, electroencephalogram (EEG), fiber photometry, and behavioral recordings. **(C)** Example image shows using Deeplabcut to track body parts. **(D)** Representative multi-recording traces during emergence from isoflurane (ISO) anesthesia from top to bottom: ISO concentration, raw EEG, EEG spectrogram, EEG total power, EEG approximate entropy, LC^NE^ activity from fiber photometry recording, and heart rate (HR). Yellow and blue dashed lines mark the start and end of the burst of increase in EEG power. Gray dashed line marks RORR. **(E)** Summary graph shows the temporal dissociation of recovery of sympathetic, cortical, subcortical, and behavioral arousal during emergence from ISO anesthesia (n=6). **(F)** Plots show averaged photometry traces with standard error of LC^NE^ activity during subcortical transition (left) and RORR (right) for 6 mice. **(G)** Plots from top to bottom show changes in EEG power, approximate entropy, LC^NE^ activity, and HR during different arousal transitions as animal is emerging from ISO anesthesia (n=6). *p<0.05. **p<0.01. n.s. not significant.

During emergence from 1.3% ISO anesthesia, the EEG increased power across a broad range of frequencies (total power 1.3±0.4×10^6^ to 6.4±1.1×10^6^ AU, p0∼.008) and exhibited a 5∼min increase in approximate entropy (20.4±3.6 to 32.6±1.3 AU, p0∼.012) before settling into stronger theta (4-8 Hz) and delta frequencies (0.5-4 Hz) reflective of the awake state (**Figure 1C**). The heart rate, as a measure of sympathetic arousal, began to increase (362±26 to 408±39 beats/min) at the EEG burst of increased power, but was not associated with increased LC^NE^ activity (Z-score -1.51±0.37 to -1.15±0.59, p0∼.256) (**Figure 1B**). Interestingly, the LC^NE^ activity had its largest tonic increase towards the end of that EEG burst (Z-score -0.83±0.62 to 4.42±1.40, p0∼.005), and a strong phasic increase at the RORR (Z-score 3.45±1.59 to 8.91±2.37, p0∼.004), signifying the return of coordinated movements (**Figure 1F**, quantified in **Figure 1G**, example raw data in **Figure 1A**, statistical analyses summarized in **Supplemental Table 1**). Additionally, our ability to dissociate sympathetic (−4.6±0.4 s relative to RORR), cortical (−2.8±0.8 s), subcortical (−0.3±1.0 s), and likely spinal activation (*i.e*. RORR) during anesthetic emergence (**Figure 1E, Figure 1B**) strongly suggests that LC^NE^ activity is tightly coupled to movement.

### LC^NE^ activity tracks in a graded manner to changes in arousal

We hypothesized that this movement-LC^NE^ coupling may be a way to tune arousal in response to movement, so we therefore characterized LC^NE^ in arousal modulation in further detail. Although LC firing has characteristic patterns associated with different stages of sleep^18^ and is known to be suppressed by intravenous anesthetics^20^, we determined whether LC^NE^ activity changes in a more binary or graded manner to changing levels of arousal.

We therefore made stepwise changes to ISO concentration from 0% to 1.5% and then back to 0% in 0.5% increments to modulate the animal’s arousal level, and observed dose-dependent changes in LC^NE^ activity (Z-score 0→ 0.25±0.29→ -0.33±0.48→ -2.0±0.7→ -2.4±0.5→ -0.29±0.08→ 1.9±0.8), corresponding changes in locomotor activity (13.9±1.0→ 26.2±4.9→ 4.5±0.9→ 0.56±0.07→ 1.7±0.2→ 5.1±1.5→ 12.6±3.7 pixels/s), and a robust neuronal hysteresis (**Figure 2A,B**). In particular, the sequential decrement in ISO dose is associated with stepwise increases in LC^NE^ activity, with each transition marked by a brief overshoot in activity (1.5% to 1.0% ISO, Z-score -3.8±0.5→ 1.4±0.8→ 0.6±0.9; 1.0% to 0.5% ISO, Z-score -1.2±0.8→ 4.0±0.5→ 2.6±0.7; 0.5% to 0% ISO, Z-score 0.08±0.48→ 4.4±0.9→ 3.4±1.1) (**Figure 2C**), reflecting strong homeostatic regulation of LC^NE^ activity. However, the overall LC^NE^ activity within 1 min of these transitions was not significantly different (**Figure 1D**). Such robust homeostatic response was also observed at RORR whether mice were emerging from 1.3% or 3.0% ISO (1.3%: Z-score -1.9±0.2→ 1.7±0.4→ 0; 3.0%: Z-score -3.6±0.7→ 2.5±0.7→ 0) (**Figure 2D-F**). Furthermore, given the temporal separation between cortical and subcortical awakening during anesthetic emergence, we sought to determine whether the increased LC^NE^ activity during RORR is transmitted to other wake-promoting centers to re-establish subcortical support of cortical wakefulness. Indeed, at RORR, the increased LC^NE^ somatic activity is correlated with increased LC^NE^ terminal activity in the medial thalamus (Thal) and basal forebrain (BF), (Z-scores LC: -1.2±0.4→ 2.5±0.6, LC^Thal^: -2.0±0.9→ 0.4±0.6; LC: -2.7±0.6→ 2.5±0.6 LC^BF^: -0.8±0.5→ 1.3±0.1), two well-established nodes in the ascending arousal pathway^21,22^ (**Figure 2G, Figure 1E,F**). Increased NE release as measured by the fluorescent NE sensor, GRAB_NE2m (Z-score Thal NE: -4.4±1.5→ 1.5±0.3; BF NE: -5.4±2.5→ 0.3±0.5), and increased postsynaptic neuronal activity (Z-score Thal: -0.1±0.5→ 3.1±0.3; BF: -0.8±0.5→ 1.3±0.2) were observed there as well. Together, these findings indicate that LC^NE^ activity strongly correlates with cortical arousal level.

**FIGURE 2.**
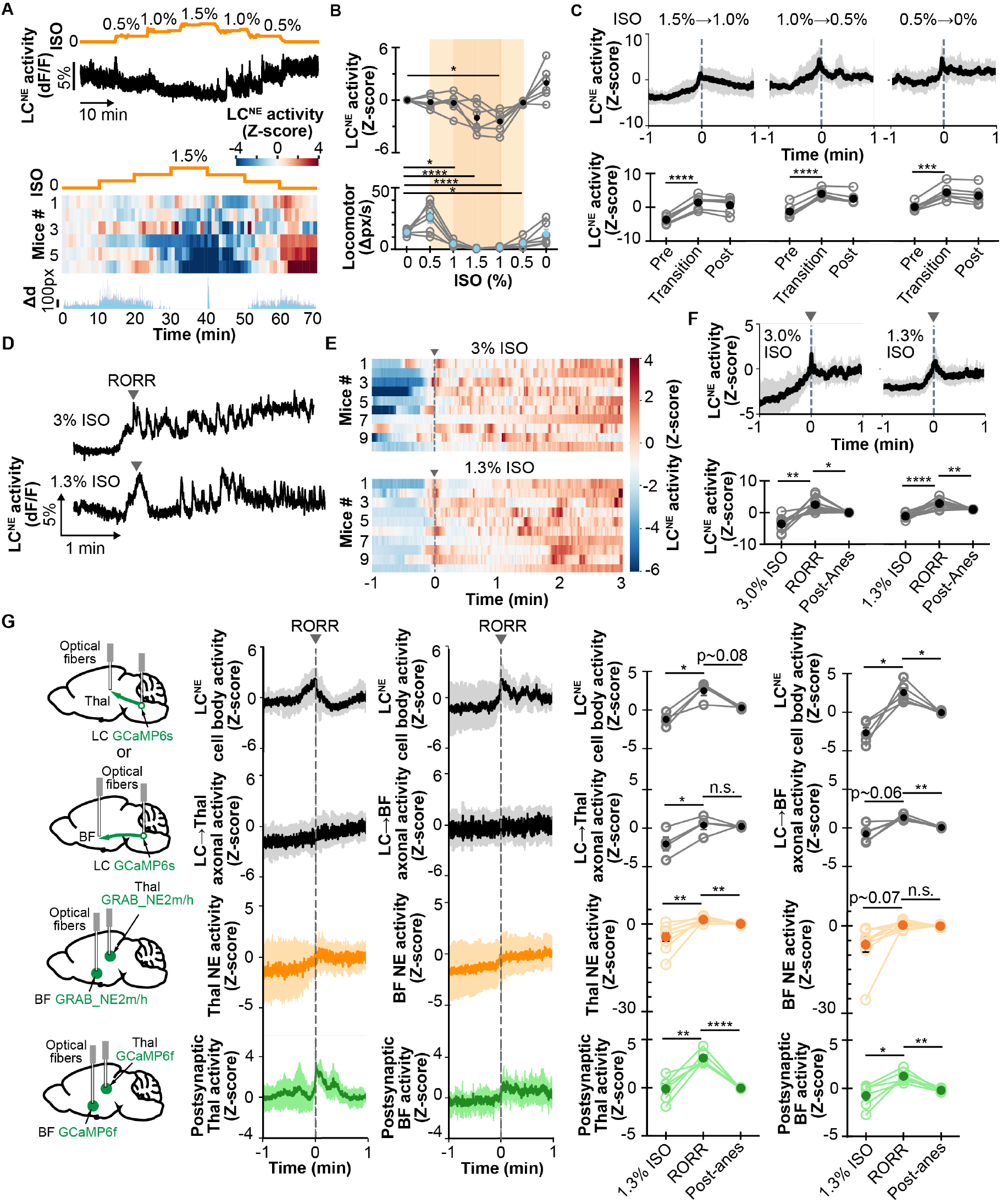
LC^NE^ activity and NE release levels correlate with changes in arousal. **(A)** Top, representative photometry trace of LC^NE^ activity during stepwise increase and decrease in isoflurane (ISO) concentration. Bottom, heatmap shows LC^NE^ activity for 6 mice for the same treatment, with corresponding averaged animal movement (Δd) from nose-tracking using Deeplabcut shown below. Increased activity seen at 40 min is an artifact from turning the mouse on its back. **(B)** Top, plot shows averaged (solid circle) and individual (open circle) mouse’s LC^NE^ activity to the different ISO concentrations. Bottom, plot shows the corresponding locomotor activity measured at different ISO concentrations. **(C)** Plots show LC^NE^ activity (top) during each stepwise decrement in ISO concentration, and its quantification (bottom) before, during, and after its stepwise increase. **(D)** Example LC^NE^ activity traces are shown during anesthetic emergence from 3% (top) or 1.3% (bottom) ISO. **(E)** Heatmaps show LC^NE^ activity during anesthetic emergence from 3% (top) or 1.3% (bottom) ISO, respectively, for 10 mice. **(F)** Plots show LC^NE^ activity (top) at RORR from 3% and 1.3% ISO and its quantification before, during, and after RORR (bottom). **(G)** Left cartoons show the expression of GCaMP6s/f to monitor neuronal activity or GRAB_NE2m/h_ to monitor NE level in either the LC, basal forebrain (BF), or medial thalamus (Thal); and the optical fiber implants. Right plots show LC^Thal^ and LC^BF^ circuit activity during RORR from 1.3% ISO anesthesia and their quantification. These include LC cell body activity and the associated LC^Thal^ (n=4) and LC^BF^ (n=5) axonal activity, Thal and BF NE (n=9/group), and Thal and BF postsynaptic activity (n=6/group). Gray arrow at time 0 marks RORR. *p<0.05. **p<0.01. ****p<0.001. n.s. not significant.

### LC^NE^ weakly promotes behavioral arousal

Despite these findings, we were still puzzled by previous studies which showed LC^NE^ to promote sleep-wake transition^6^ and anesthetic emergence, but to be relatively ineffective in promoting reanimation under anesthesia^23^. Therefore, we next determined whether the LC^NE^ system is necessary to promote wakefulness by investigating its specific causal role in arousal modulation using a series of selective pharmacological manipulations which target various presynaptic actions of LC^NE^ neuromodulation. Specifically, we used dexmedetomidine (DEXMED, α_2_-AR agonist), atipamezole (ATI, α_2_-AR antagonist), and atomoxetine (ATOM, NE transporter inhibitor); and compared LC^NE^ activity during emergence from ISO anesthesia and emergence time to those of saline control (SAL) (**Figure 3A**). While the LC^NE^ phasic firing at RORR (Z-scores SAL: -0.2±0.3→ 4.5±0.8, DEXMED: -0.8±0.5→ 4.9±1.5, ATI: 0.1±0.3→ 4.6±0.6, ATOM: -0.5±0.3→ 2.5±0.6) and emergence time (SAL: 103±22 s, DEXMED: 157±48 s, ATI: 83±20 s, ATOM: 94±13 s) did not differ between treatments (**Figure 3A,B**), the LC^NE^ calcium activity pattern did however exhibit some subtle differences. For instance, for SAL and ATI treatments, the LC^NE^ activity increased post-anesthesia compared to preanesthesia, but did not differ in DEXMED and ATOM treatments (ΔZ-score SAL: 1.7±0.5, DEXMED: 0.7±0.7, ATI: 1.0±0.4, ATOM: -0.3±0.3) (**Figure 3C**). Combined with the observation that LC^NE^ activity at RORR trended higher for DEXMED compared to SAL (ΔZ-score SAL: 4.7±0.8, DEXMED: 5.7±1.2, ATI: 4.4±0.7, ATOM: 3.0±0.8) (**Figure 3D**), these results suggest that the LC^NE^ homeostatic response remains intact with DEXMED treatment, even though its overall activity is suppressed by the selective α_2_ agonism. By contrast, LC^NE^ activity at RORR increased significantly less after ATOM treatment, suggesting a possible ceiling effect that limits both the LC^NE^ homeostatic response and its tonic activity. Overall, these pharmacological studies support a relatively weak role of LC^NE^ in promoting behavioral arousal at the lower end of the arousal spectrum.

**FIGURE 3.**
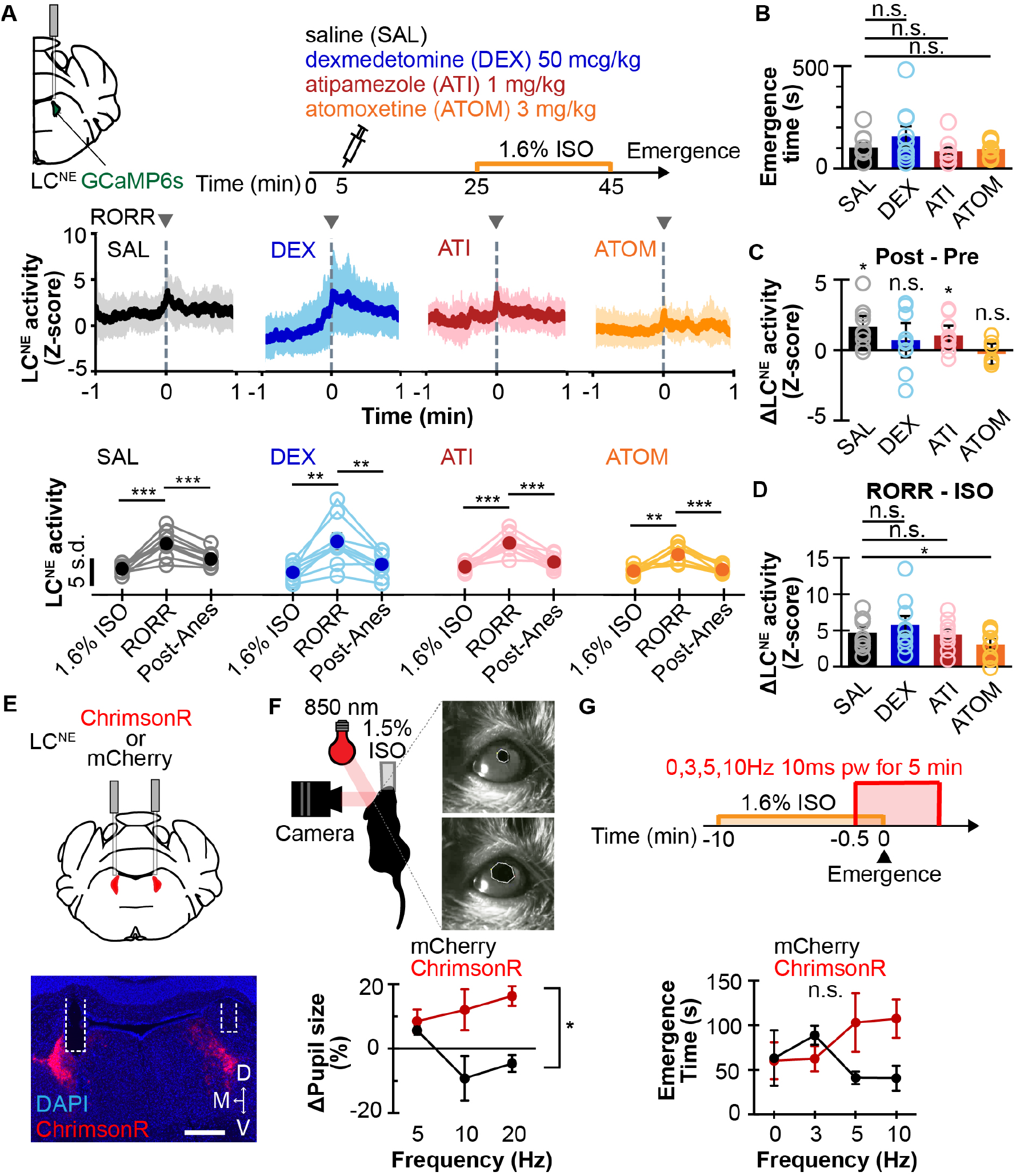
Effect of LC^NE^ pharmacological modulation on LC^NE^ during emergence events. **(A)** Top, diagrams show GCaMP6s expression to monitor LC^NE^ activity and experimental design to modulate LC^NE^ circuits pharmacologically using saline (SAL), dexmedetomidine (DEX), atipamezole (ATI), and atomoxetine (ATOM) during emergence from 1.3% isoflurane (ISO) anesthesia. Averaged traces of LC^NE^ activity at RORR are shown for the four treatments (middle), with the corresponding quantification (bottom). **(B)** Emergence time (*i.e*. time to RORR) is shown for the four different drug treatments. **(C)** Changes in tonic LC^NE^ level from before to after RORR are shown for the four different drug treatments. **(D)** Changes in LC^NE^ level from anesthetized to RORR are shown for the four different drug treatments. **(E)** Top, schematic shows bilateral expression of ChrimsonR or mCherry control, and bilateral optical fiber implant, in the LC^NE^ of DBH-cre mice. Bottom, example histology of ChrimsonR expression and optical fiber tracks (white dashed line) in the LC^NE^. Scale bar is 500 μm. **(F)** Top, schematic shows anesthetized mouse (at 1.5% ISO) in pupillometry setup (left) and closeups of changing pupil sizes as tracked by Deeplabcut. Bottom, plot shows changes in pupil size in response to LC^NE^ photostimulation in animals expressing either ChrimsonR (n=4) or mCherry (n=3). **(G)** Top, diagram shows experimental design for LC^NE^ photoactivation during emergence from 1.6% ISO. Bottom, plot shows emergence time in ChrimsonR- (n=5) and mCherry-expressing (n=3) mice. *p<0.05, **p<0.01, ****p<0.001, n.s. not significant.

### LC^NE^ potently activates sympathetic arousal

Besides promoting cortical arousal, LC^NE^ activation induces sympathetic arousal, even under anesthesia^24^. To examine this more specifically, we photoactivated the LC^NE^ neurons expressing ChrimsonR (optogenetic actuator) under 1.5% ISO, and found a dose-dependent pupil dilation not seen in mCherry controls (Δpupil size: ChrimsonR, 5 Hz: 8.5%±3.6%, 10 Hz: 12.0%±6.3%, 20 Hz: 16.3%±3.1%; mCherry, 5 Hz: 5.6%±1.3%, 10 Hz -9.2%±6.9%, 20 Hz: -4.6%±2.6%) (**Figure 3E,F**), consistent with sympathetic activation. Unexpectedly, we did not find any significant difference in emergence time when the LC^NE^ neurons were photoactivated during emergence from 1.6% ISO (ChrimsonR, 0 Hz: 60±21 s, 3 Hz: 63±14 s, 5 Hz: 103±33 s, 10 Hz: 107±21 s; mCherry, 0 Hz: 63±31 s, 3 Hz: 89±11 s, 5 Hz: 41±7 s; 10 Hz: 41±14 s) (**Figure 3G**). Additionally, we retrogradely expressed inhibitory designer receptors (Gi DREADD) in the LC-medial thalamic and LC-basal forebrain circuits. Interestingly, compared to the saline control, administrating clozapine-N-oxide (CNO) to activate Gi DREADD to inhibit these two circuits did not significantly alter emergence time from ISO anesthesia (**Figure 1G**). Taken together, whereas the photometry recording of LC^NE^ calcium dynamics show behavioral arousal coupled to LC^NE^ activation, the results from the pharmacological, optogenetic, and chemogenetic modulation of LC^NE^ demonstrate overall a limited causal role of LC^NE^ in promoting behavioral arousal.

### LC^NE^ activity is tuned to changes in movements

Although our anesthetic emergence model shows a robust movement-to-LC^NE^ coupling link at RORR, it is unclear whether this coupling is important during natural conditions. Therefore, we sought to examine the relationship between movement and LC^NE^ activity in awake freely-moving mice. Using GCaMP6s-expressing LC^NE^ neurons to monitor activity and Deeplabcut^25^ to track mouse movement kinematics, we found that movement initiation and arrest events are associated with distinct patterns of LC^NE^ activity. In particular, while movement initiations coincide with a dip followed by an increase in LC^NE^ activity (Z-scores 0.11±0.05→ -0.31±0.06→ 0.35±0.06→ 0.17±0.04), movement arrests show a gradual ramp in activity followed by a decrease (Z-scores 0.04±0.01→ 0.22±0.04→ -0.38±0.11→ 0.10±0.03) (**Figure 4A**). These changes were not movement artifacts as they were not observed in the control 405nm isosbestic channel (**Figure 3A**). By averaging the LC^NE^ activity at movement initiations, we generated a computational template model that captures the biphasic response (**Figure 4B,C, Figure 3B**). Sliding this model along a photometry trace while measuring the cross-correlational coefficient, we were able to optimize the combination of local maxima and thresholding to detect a large proportion of movement initiation LC^NE^ activity events, including very subtle movements such as a brief head re-orientation (**Figure 4D,E**). When these detected events were averaged, they recapitulate the template as expected, but also revealed the specific mouse movement transitions (**Figure 2F**). Moreover, these events show a distribution of activity that is slightly offset compared to the actual movement initiations, suggesting that LC^NE^ activation follows the movement initiations (**Figure 2G**). Similar results are also found for movement arrests, though the template tends to fit less well when compared to that for initiations, as reflected by the overall lower ceiling in the true positive rate (**Figure 4H-L**). In addition, when compared to baseline, SAL, DEXMED, ATI, and ATOM treatment did not alter the LC^NE^ patterns associated with movement initiations and arrests (**Figure 3C,D**). Together, these template-based model predictions suggest that stereotyped LC^NE^ activity is tightly coupled to movements in awake animals.

**FIGURE 4.**
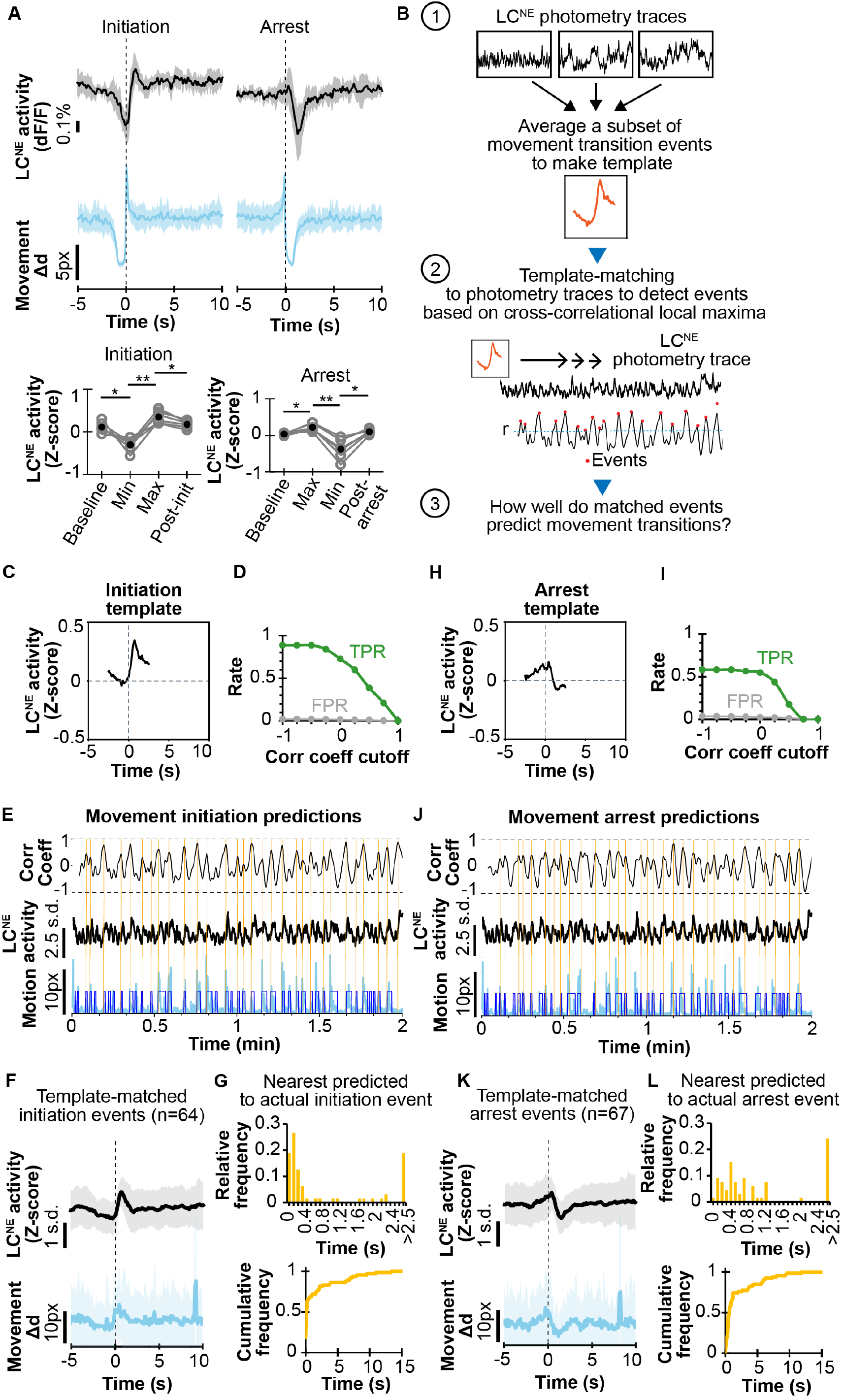
Modeling Movement-LC coupling in awake animals. **(A)** Top, plots show the corresponding LC^NE^ activity from fiber photometry recording in response to movement initiation or arrest, as tracked by Deeplabcut, for 6 mice. Bottom, quantification of changes in LC^NE^ activity during these movement transitions. Traces are averaged across events and mice. *p<0.05. **p<0.01. **(B)** Flow diagram shows the steps from generating template, to using template-matching cross-correlational-based detection of neuronal activity events, and finally to investigate how predictive are these events for movement transitions. **(C)** Plot shows template of LC^NE^ activity associated with movement initiation generated from averaging events across 9 mice. **(D)** Plot shows the true positive rate (TPR) and false positive rate (FPR) for the example mouse using various correlation coefficient cutoff when performing template matching for movement initiation. **(E)** An example of events detected from matching LC^NE^ activity template in **(C)** to photometry trace (middle), based on local maxima of rolling correlation coefficient (top), marked as orange lines and compared to the true binary movement categorization (dark blue) overlying the raw movement trace (bottom). **(F)** Averaged movement trace (bottom) associated with template detected events (n=64) in **(E)**. Averaged LC^NE^ activity for these events is shown on top. **(G)** Top plot shows relative frequency of the time difference between the template detected events and nearest true movement initiations. Bottom plot shows the cumulative frequency of these time differences. **(H)** Plot shows template of LC^NE^ activity associated with movement arrest generated from averaging events across 9 mice. **(I)** Plot shows TPR and FPR for using various correlation coefficient cutoff when performing template matching for movement arrest. **(J)** Example of events detected from matching LC^NE^ activity template in **(H)** to photometry trace (middle), based on local maxima of rolling correlation coefficient (top), marked as orange lines and compared to the true binary movement categorization (dark blue) overlying the raw movement trace (bottom). **(K)** Averaged movement trace (bottom) associated with template detected events (n=67) in **(J)**. Averaged LC^NE^ activity for these events is shown on top. **(L)** Top plot shows relative frequency of the time difference between the template detected events and nearest true movement arrests. Bottom plot shows the cumulative frequency of these time differences. *p<0.05, **p<0.01, ***p<0.001, n.s. not significant.

We also examined the movement-LC^NE^ coupling before and after ISO anesthesia by looking at LC^NE^ patterns during movement transitions prior and for up to 90 min after exposure to 10 min of 1.3% ISO anesthesia (**Figure 4A**). Interestingly, we also observed a noteworthy delay in the characteristic LC^NE^ pattern for both movement initiations and arrests in a subset of animals after anesthesia, despite appearing to have grossly intact movements (**Figure 4B-D**). These observations indicate a coupling between movement and LC^NE^ that is not causal, but likely is regulated by unique upstream inputs and/or recurrent cortico-LC loops which have been postulated to exist. They also suggest that anesthetic disruption of specific brain circuits may persist even when the animal appears behaviorally to have “recovered” from anesthesia. Thus, further studies are warranted to explore the clinical implications of these findings. In total, these results with our template model demonstrates a stereotypic pattern in movement-LC^NE^ coupling that may be useful in detecting subtle brain dysfunction.

## 3 DISCUSSION

Using emergence from ISO as a tool to dissociate various arousal components, we report here that significant increases in LC^NE^ activity occur after cortical and sympathetic activation, suggesting that LC^NE^ is not a major contributor to wakefulness at the lower end of the arousal spectrum. Indeed, this is consistent with prior research that showed stimulation of the ventral tegmental area dopaminergic neurons is capable of inducing reanimation under ISO anesthesia^26^, while increasing LC^NE^ activity using Gq-activating designer receptors could not^23^. Furthermore, although chemogenetic activation of LC^NE^ has been shown to facilitate anesthetic emergence, we did not observe the same effect with optogenetic activation, possibly due to limited window of NE increase before its depletion^27^ or the limited ability of the opsins to robustly mobilize intracellular calcium stores. Although loss of righting has been well correlated with the loss of consciousness during anesthetic induction^28^, it is often assumed that RORR also correlates with the return of consciousness. However, given the observed cephalad-caudal progression of central nervous system activation during ISO emergence, it is possible that consciousness actually returns prior to the expression of RORR behavior.

During ISO induction, locomotor activity dramatically decreased at 1.0% ISO, whereas LC^NE^ activity exhibited a steep decrease at 1.5% ISO (**Figure 2A,B**), suggesting that anesthesia may dissociate movement-LC^NE^ coupling. This is also consistent with the variability in the recovery of this coupling after ISO anesthesia (**Figure 4**). Moreover, the observed hysteresis of LC^NE^ activity between anesthetic induction and emergence is consistent with the proposed neuronal inertia as underlying the anesthetic hysteresis observed at the organismal level^29^.

In both rapid-eye movement (REM) sleep and cataplexy in which there is a loss of muscle tone, LC remains largely silent^14,30^. A more recent work showed that activation of α_1_-ARs during a cataplexy episode increased postural muscle tone, and concluded that the noradrenergic system couples motor and arousal systems^15^. This is further supported anatomically by the direct LC projections to the motor cortices^31^. However, our results did not establish a strong causal role of LC^NE^ in promoting movement, but rather, movement changes associate with a coordinated pattern of LC^NE^ activity. How this coordination occurs is not entirely clear, as it is not known whether motor cortices directly project to the LC^NE^. The fact that in certain mice after anesthesia, the movement-LC^NE^ coupling can be delayed on the orders of hundreds of seconds seems to suggest that secondary upstream regulators likely act to coordinate movement and LC^NE^ activity. While it is unclear whether LC^NE^ is responding to movement *per se* rather than to muscle stretch, this movement-LC^NE^ coupling identified in this study offers a possible mechanism for afferentation theory in which muscle afferent activity is thought to activate key brain arousal centers^32^.

Assessing movement-LC^NE^ coupling has potential translational utility. The observed differential recovery in movement-LC^NE^ coupling after anesthesia may be an avenue for examining perioperative neurocognitive disorders such as delirium and impaired cognition^33^. Additionally, neurodegenerative diseases such as Alzheimer’s and Parkinson’s involve the loss of LC^NE^ neurons, and the impaired movement-LC^NE^ coupling may be a major consequence that can explain pathophysiological features such as dysautonomia and impaired movements^34,35^. Movement-LC^NE^ coupling may also be important in REM sleep disorders in which muscle tone may be inappropriately lost or preserved^36^. Thus, further studies are warranted to investigate the roles of LC^NE^ neuronal activity in movement initiations and arrests under pathological states in relevant disease animal models.

In summary, we report here a robust movement-LC^NE^ coupling, which together with a nuanced role of LC^NE^ in promoting cortical and sympathetic arousal, suggest that LC^NE^ activity plays an important role in tuning arousal to specific movement events. The movement-LC^NE^ coupling may also provide translational insights into pathophysiology in neurodegenerative diseases, approach/avoidance, motivated behavior, sleep disorders, and perioperative cognitive disorders.

## Abbreviations

LC: locus coeruleus
NE: norepinephrine

## 4 METHODS

### Animals and surgery

Adult male and female Dbh-cre C57BL/6J mice (Dbh^*tm3.2(cre)Pjen*^, Jackson Laboratory #033951) were group housed on a 12:12 h reverse light-dark cycle and given food and water *ad libitum*. Surgeries were performed on mice ages >8 weeks under 1.5-2.0% isoflurane. Mice were injected in the LC (AP -5.45mm, ML +1.1mm, DV -4.2 to -3.7mm DV to bregma) using a medial-facing beveled 1 µl Hamilton syringe. A blunt tip 1 µl Hamilton syringe was used for injections to the posterior basal forebrain (BF) (AP +0.1mm, ML +1.3mm, DV -5.3 to -5.15mm) and the medial thalamus (mThal) (AP -1.7mm, ML +1.0mm at 15° angle, DV -3.95 to -3.8mm). Respective 400 µm optical fibers (Doric Inc., MFC_400/430-0.48_MF2.5_FLT) were placed along the same AP and ML coordinates except DV is -3.8mm for LC, -5.2mm for BF, and -3.8mm for mThal for fiber photometry; and secured using Metabond (Parkell #S380). For LC photoactivation, 200 µm optical fibers were implanted at (AP -5.45mm, ML +/-0.9mm, DV -3.8mm). Mice recovered for at least 4 weeks before experiments to allow optimal viral expression. All experiments are in accordance to IACUC protocol.

### Histology

Briefly, mice were euthanized with sodium pentobarbital and transcardially perfused with 4% paraformaldehyde (PFA), post-fixed for 1-3 days in 4% PFA, and then cryo-protected in 30% sucrose. Brain sections (30-100µm) were collected and kept in 0.1M phosphate buffer at 4° C until mounting. Sections were mounted with VectaShield Vibrance Hardset mounting medium (Vector Laboratories) with DAPI, and coverslips placed. Images were acquired on an epifluorescent microscope (Leica DFC700T).

### Viruses

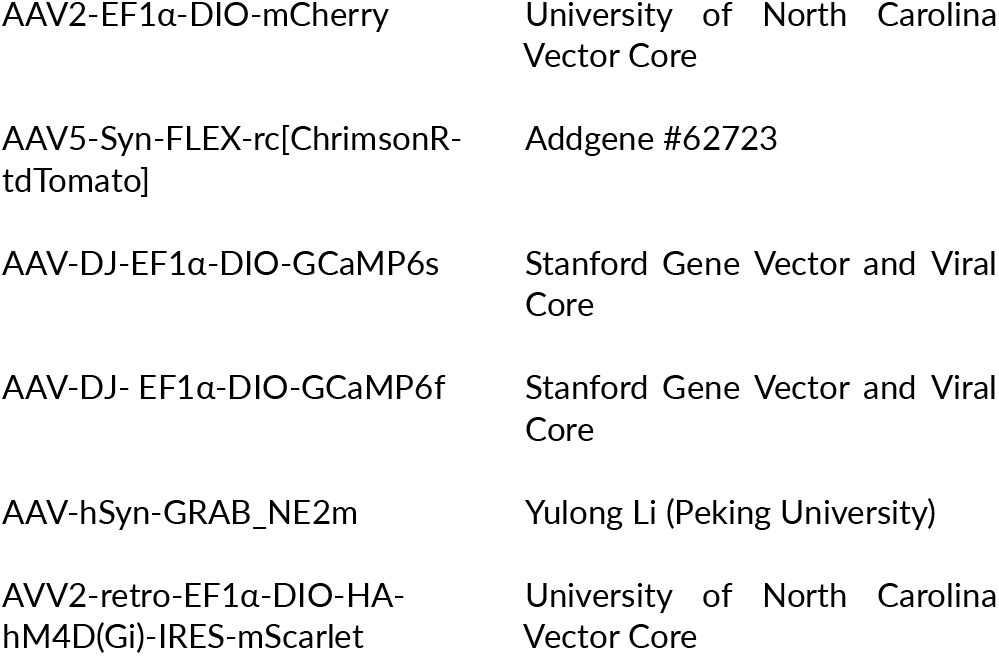

|  |  |
| --- | --- |
| AAV2-EF1α-DIO-mCherry | University of North Carolina Vector Core |
| AAV5-Syn-FLEX-rc[ChrimsonR-tdTomato] | Addgene #62723 |
| AAV-DJ-EF1α-DIO-GCaMP6s | Stanford Gene Vector and Viral Core |
| AAV-DJ-EF1α-DIO-GCaMP6f | Stanford Gene Vector and Viral Core |
| AAV-hSyn-GRAB_NE2m | Yulong Li (Peking University) |
| AAV2-retro-EF1α-DIO-HA-hM4D(Gi)-IRES-mScarlet | University of North Carolina Vector Core |

### Fiber photometry and photostimulation

Fiber photometry recordings of the genetically encoded fluorescent sensors were performed as previously described^37^. Briefly, the implanted fiber was connected to the patch cable via a ferrule sleeve. A real-time processor (TDT RZ5P, sampling rate of 1017.25Hz) recorded the filtered emission (Doric FMC4 filter cube) from the fluorescent sensor at upon excitation at 470nm and 405nm using the accompanied TDT Synapse software. For terminal stimulations, the same photometry fiber served as conduit for stimulation whereas a dual commutator was used for bilateral LC Photostimulation. In both cases, 625nm LED was adjusted to a power output of 2∼.5-5 mW/mm^2^ at the fiber tip.

Photometry recordings were analyzed using custom python script as before^27^. Baseline drift from photobleaching artifact was corrected with an exponential decay curve. For GRAB_NE and GCaMP signals, dF/F was calculated as the linear least squared fit of 405nm signals subtracted from the 470nm signals. Z-scores were calculated using the mean and standard deviation of a 5-10 min baseline before photostimulation.

### Electroencephalogram (EEG)

Mice were implanted with 2×2 pinbrick in the left frontal brain region (+1.2 AP, -0.65 ML), (+2.5 AP, -0.65 ML), (+2.5 AP, -1.95 ML) using a modified protocol from previously described^38^. Briefly, the pinbrick is attached to an Intan headstage, which is connected to the Open Ephys acquisition board. Ground and reference Intan channels were wired together. Associated Open Ephys GUI was used for data acquisition. Analysis was performed using custom python script.

### Vitals monitoring

Vitals (heart rate, respiratory rate, oxygen saturation, and pulse distension) were acquired via the collar sensor in both anesthetized and awake animals using the MouseOx system.

### Pupillometry

Protocol is based on our prior work^39^. Mice were placed in a nosecone and anesthetized with 1.0-1.5% ISO. In a dimly lit room, a FLIR camera (FFY-U3-16S2M-S) and a far-infrared lamp were directed at the right eye. Spinnaker software was used for recording. Photostimulation was performed as described above.

### Anesthesia experiments

Mice was placed in a custom cylindrical chamber (10 in diameter x 10 in height). The lid contains a slit through which cables may pass. ISO was delivered via an inlet on the bottom of the cylindrical chamber while vacuum was connected to an outlet on top. A small tubing near the bottom of the chamber was used for gas sampling that is fed to a gas analyzer (Ohmeda 5250 RGM, GE Healthcare, Madison, WI). Heating pad was positioned at the bottom of the cylinder. For experiments measuring emergence time, anesthetized mice were immediately transferred to an adjacent open container and placed on its back, and the return of righting is marked by when all 4 paws have touched the container floor.

### Behavioral analysis

Mice behavior was video recorded. The nose, ears, and tail base of a small subset of images were manually labeled and used in the Deeplab-cut pipeline^40^. Frame-to-frame displacement was calculated via custom python script. Template generation for movement initiation and arrest used a pause interval of ≥1 s and movement threshold <10 pixel to denote period without movement. For movement initiation, template is bounded by 1.5 s before and 0.7 s after event while for movement arrest, template is bounded by 1 s before 1.6 s after event. These were chosen to minimize template length while capturing the distinct patterns of movement transitions. The template is then slid along the photometry trace while the Pearson correlation coefficient (r) is calculated for each position, and local peaks with r>0 are denoted as events. These events are aligned to movement for calculating the true and false positive rates.

### Drugs and treatments

All drugs were administered intraperitoneally (*i.p*.). These include dexmedetomidine (100 mcg/ml Piramal Critical Care, PSLAB-020872-00) diluted to 5 mcg/ml in saline, atipazemole made to 0.1 mg/ml in saline, atomoxetine hydrochloride (Thermoscientific, Cat 467680010) make to 0.3mg/ml in saline. These drugs were administered at least 15-20 min prior to experiments unless otherwise noted. For DREADD experiments, clozapine N-oxide (CNO) (5 mg/kg, Enzo Life Sciences, Cat#BML-NS105) was administered at least 30 minutes prior to anesthetic emergence.

### Statistical analysis

All summary data are expressed as mean±SEM. Statistical significance was denoted as *p<0.05, **p<0.01, ***p<0.001, ****p<0.001 as determined by Student’s t-test, one-way or two-way repeated measure analysis of variance (ANOVA), followed by either Tukey’s or Dunnett’s *post-hoc* test. Statistical tests were performed in Excel, R, and Prism. A summary of the statistical tests performed and the associated p-values are provided in **Supplemental Table 1**.

## DATA AND CODE AVAILABILITY

Custom python analysis code, and photometry data will be provided upon request.

## AUTHOR CONTRIBUTIONS

L.L. A.R. E.M.L. and M.O.T. collected data. L.L. and M.R.B. provided resources, designed experiments, analyzed data, and wrote the paper.

## ACKNOWLEDGMENTS

We thank all of Bruchas lab for their helpful insights and discussions with manuscript. We thank Yulong Li (Peking University) for providing GRAB_NE viruses. We acknowledge our funding sources; FAER MRTG (L.L.), K99/R00 DA053336 (L.L.), Mary Gates Research Scholarship (E.M.L.), and R01 MH112355 (M.R.B).

---

**Supplemental Fig 1.**
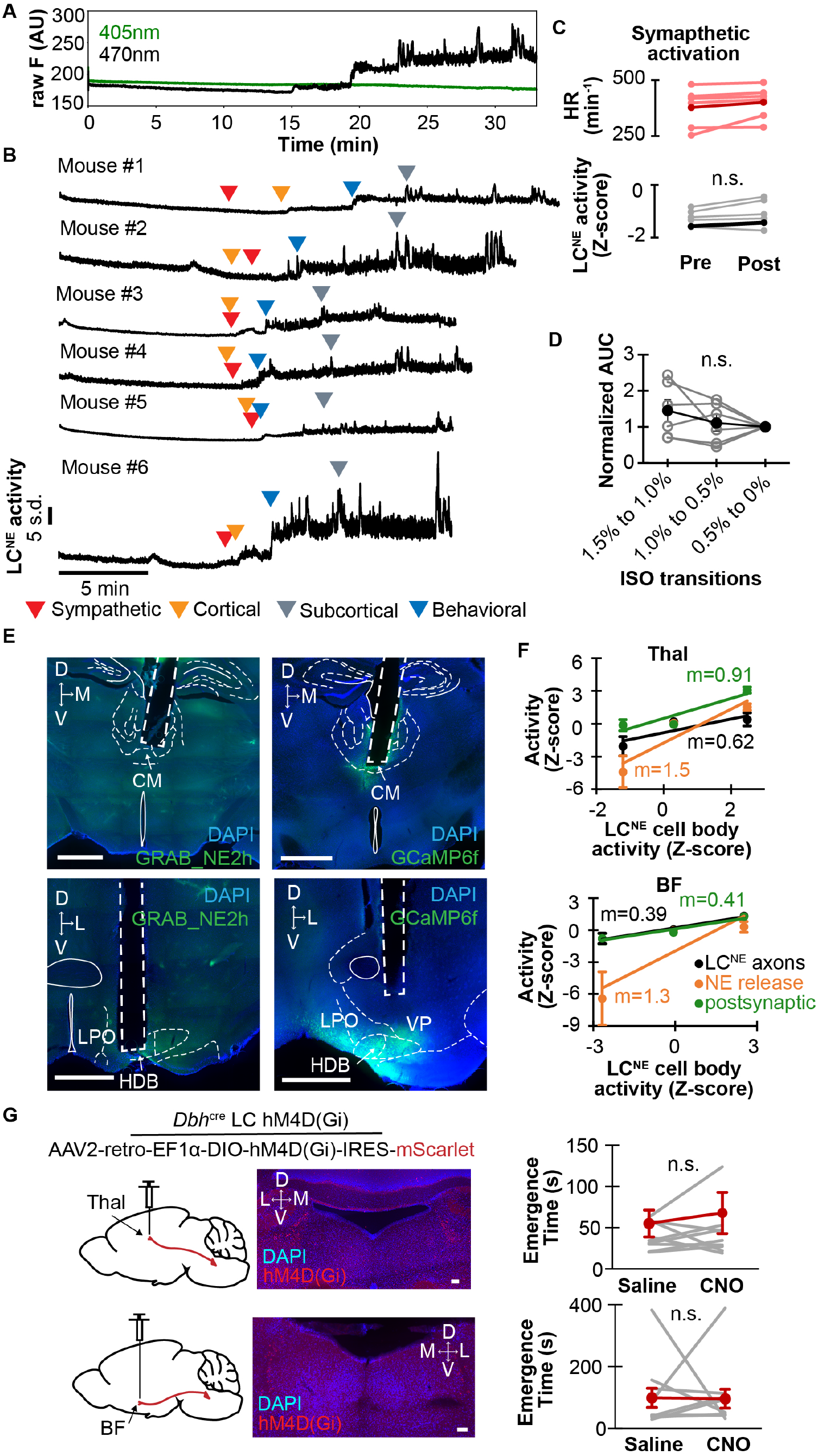
Role of LC^NE^ during anesthetic emergence. **(A)** Representative raw photometry traces at 405nm (isosbestic) and 470nm of GCaMP6s-expressing LC^NE^ during emergence from isoflurane (ISO) anesthesia. **(B)** Photometry traces of LC^NE^ activity for individual mouse during emergence from 1.3% ISO, with arrows indicating sympathetic, cortical, subcortical, and behavioral arousal transitions. **(C)** LC^NE^ activity (bottom, Z-scores-1.6±0.4→ -1.4±0.4, p 0.178) during sympathetic arousal transitions (top, 378±36 beats/min→ 401±30 beats/min), as reflected by increased heart rate (HR). **(D)** Normalized area under curve (AUC) for LC^NE^ activity during 1 min after the stepped increase in response to stepwise decrement in ISO concentration (1.5% to 1.0%: 1.5±0.3, 1.0% to 0.5%: 1.1±0.2, 0.5% to 0%: 1±0). **(E)** Example of mice histology showing on the left, GRAB_NE2h_ expression and optical placement in the medial thalamus (Thal), top; and in the posterior basal forebrain (BF), bottom; and on the right, GCaMP6f expression and optical placement in Thal, top; and in BF, bottom. **(F)** Plots show relation between LC^NE^ cell body activity at RORR and NE release and postsynaptic activity in Thal (top) and BF (bottom) **(G)** Left schematics show injection of a Gi-DREADD expressing retrograde virus in the medial thalamus (top) and BF (bottom) in DBH-cre mice to selectively target LC-Thal and LC-BF circuits, respectively. Middle histology images show the respective Gi-DREADD expressions in the LC. Right graphs show the emergence time of these respective mice injected intraperitoneally with saline or 5 mg/kg CNO at least 30 min before emerging from 10 min of 1.6% ISO (LC^Thal^: SAL 55±16 s, CNO: 68±25 s, n=10; LC^BF^: SAL: 98±31 s, CNO: 95±31 s, n=10). n.s. not significant.

**Supplemental Fig 2.**
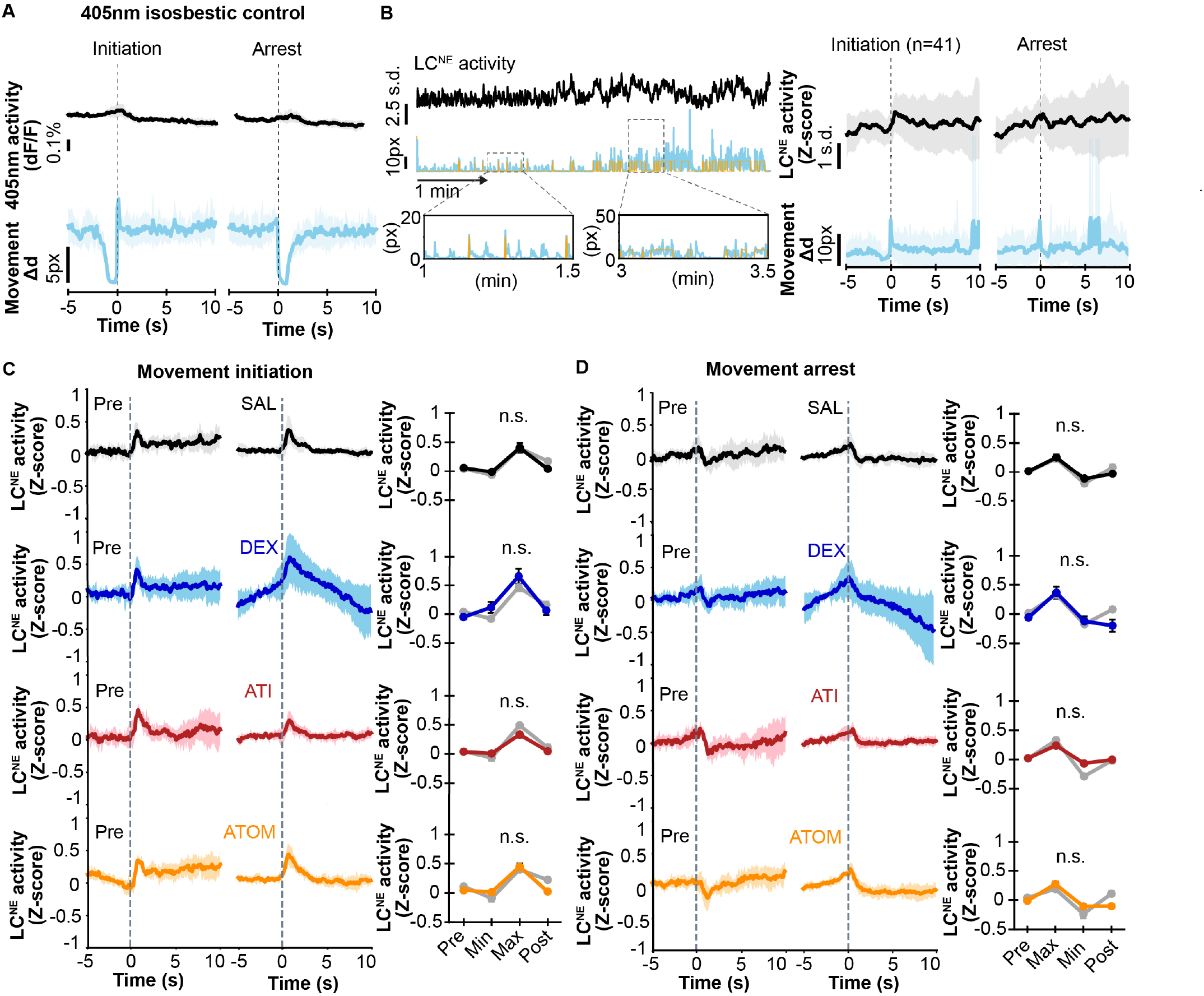
Association of movement and LC^NE^ activity. **(A)** Photometry traces of 405nm (isosbestic) channel for LC^NE^ activity (top) associated with movement transitions (left, initiation; right, arrest) are shown (n=6 mice). **(B)** Representative example of event discovery for generating LC^NE^ activity template. Top, LC^NE^ activity trace (black) and movement trace (light blue), with the overlying binary categorization of movement (orange) to demarcate events for analysis. Zoomed-in boxes provide two closeup examples. Bottom, patterns of averaged LC^NE^ activity (black) associated with n=41 movement initiation events (left, light blue) and movement arrest events (right, light blue) for an example mouse. **(C)** Left plots show LC^NE^ activity associated with movement initiations in awake mice before and after treatment with saline control (SAL), dexmedetomidine (DEX), atipamezole (ATI), and atomoxetine (ATOM). Right plots are quantifications of the corresponding patterns of LC^NE^ activity, with the before treatment in gray and after treatment in color. **(D)** Same as in **(A)** except associated with movement arrests.

**Supplemental Fig 3.**
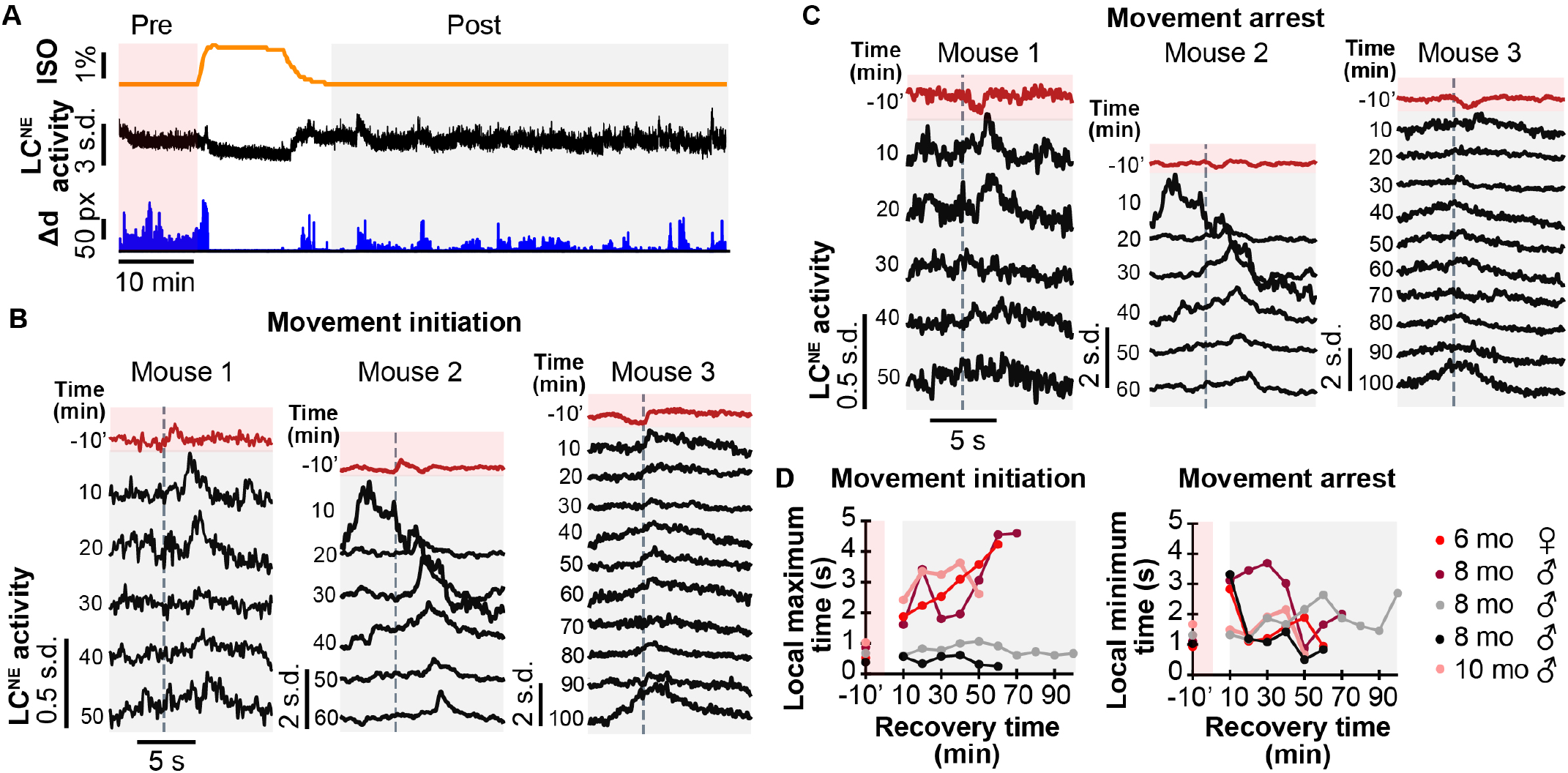
Variations in movement-LC^NE^ coupling after anesthesia. **(A)** Representative anesthetic experiment, with 10-15 min treatment of 1.3% isoflurane (ISO), followed by washout (top), and the corresponding LC^NE^ activity (middle) and movement (bottom). Preanesthetic baseline for analysis is shaded red whereas postanesthetic window of analysis is shaded gray. **(B)** Examples of LC^NE^ activity associated with movement initiations at baseline and during anesthetic recovery for 3 mice. Mouse 1 and 2 show delayed movement-LC coupling while mouse 3 does not. **(C)** Same as **(B)** except showing activity associated with movement arrests. **(D)** Plots show time to local maxima of LC^NE^ activity at movement initiations (left) and time to local minima of LC^NE^ activity at movement arrests (right) pre- and post-anesthetic treatment.

